# Metabolome and transcriptome profiling of root chicory provide insights into laticifer development and specialized metabolism

**DOI:** 10.1101/2024.01.02.573856

**Authors:** Khabat Vahabi, Gerd U. Balcke, Johanna C. Hakkert, Ingrid M. van der Meer, Benedikt Athmer, Alain Tissier

## Abstract

Chicory roots produce inulin, a dietary fiber, as well as large quantities of bitter sesquiterpene lactones (STLs), which have valuable biological activities. In an effort to understand the compartmentalization of metabolism within chicory roots and the molecular basis of the development of laticifers that produce the chicory latex, we performed metabolomics and transcriptomics profiling. GC-MS and LC-MS identified a total of 22 580 features of which 135 were differentially abundant between cell types. Further analysis indicated that the major STLs accumulated primarily in the latex. Gene expression of known STL pathway genes indicates a compartmentalization of the biosynthesis across multiple tissues, with implications regarding the trafficking of pathway intermediates. Phytohormone measurements and gene expression analysis point to a major role for jasmonate signaling in the development and differentiation of laticifers. Furthermore, inulin accumulates mostly outside the laticifers but expression of inulin metabolic genes also point to a complex distribution and trafficking of inulin or inulin precursors across different root compartments. Altogether, the data presented here constitute a unique resource to investigate several biological processes in chicory roots, including laticifer development, STL biosynthesis and transport and inulin biosynthesis regulation.

**Significance statement:** A combination of transcriptomics, targeted and untargeted metabolomics of different tissues of chicory roots was generated. These data constitute a resource basis for the investigation of various processes taking place in chicory taproots, including sesquiterpene lactone biosynthesis, laticifer development and inulin biosynthesis and trafficking.

## Introduction

Root chicory (*Cichorium intybus* var. *sativum* L.) is a variety of common chicory that is grown for its inulin-containing roots. Inulin is a soluble fructan polymer with a terminating glucose residue that is used as a dietary fiber and sugar replacement in many food products, such as yogurts, other dairy products and bakery goods. According to the Food and Agriculture Organization (FAO) Gross Production Value (current million US$) of root chicory is about 555.85 million dollars per year (**Fig. S1A**). Most of the production of root chicory comes from Europe (**Fig. S1B**), with Belgium, France, the Netherlands and Poland ranked as the main producers (**Fig. S1C**).

A traditional use of root chicory that goes back to the 18^th^ and 19^th^ centuries in Germany and France is as coffee replacement. Although chicory does not contain any caffeine, it has bitter compounds that confer a similar taste to coffee. These bitter compounds are sesquiterpene lactones (STLs), which occur frequently in the Asteraceae family. In many plants of the Asteraceae family, the STLs accumulate to high levels in glandular trichomes or in latex, which is a secretion of the laticifer cells. Chicory roots produce an abundant latex containing the STLs that are derived from germacrene A and include 8-deoxylactucin (DL), lactucin, lactucopicrin and their oxalate conjugates (Kraker *et al.*, 1998) (**Fig. 1**). STLs constitute a large and diverse group of sesquiterpenoids, with various pharmacological and biological activities, including antibacterial, anti-inflammatory and anti-helminthic (Li *et al.*, 2018, Woolsey *et al.*, 2019). Chicory STLs also possess anti-helminthic and anti-inflammatory activities (Peña-Espinoza *et al.*, 2015, Matos *et al.*, 2020, Peña-Espinoza *et al.*, 2020, Häkkinen *et al.*, 2021). Probably the most famous of STLs is artemisinin, a compound with anti-malarial activity produced in the glandular trichomes of sweet wormwood (*Artemisia annua*) (Xiao *et al.*, 2016). In only few cases is the function of STLs in their natural environment known. One example is in dandelion (*Taraxacum officinale* agg.), where a STL produced in latex, a glycosylated taraxinic acid, confers resistance to insect larvae that feed on the roots (Huber *et al.*, 2016a, Huber *et al.*, 2016b).

STLs interfere with the extraction of inulin from chicory roots. Therefore, there is interest in generating chicory lines with reduced STL content. A recent report describes the generation of clustered regularly interspaced short palindromic repeats associated protein 9 (CRISPR/Cas9) knock-outs of chicory genes encoding germacrene A synthase (GAS) (Cankar *et al.*, 2021). All four copies of the chicory *GAS* genes were disrupted, leading to a complete loss of STL production. Interestingly this was accompanied by increased accumulation of squalene, which derives from the same precursor as the STLs, namely farnesyl diphosphate. On the other hand, because of their potential health benefits and various biological activities, STLs from chicory may also constitute a breeding target, for example to modify the STL content for the accumulation of specific compounds or to globally increase STL productivity in roots. In related *Lactuca* species and in chicory it was shown that STLs are present in the latex (Sessa *et al.*, 2000). In different plant species, the latex contains high concentrations of various secondary metabolites and plays a protective role against microbes, pests and wounding by accumulating high levels of toxic proteins and metabolites (Makita *et al.*, 2017, Kitajima *et al.*, 2018). Thus, knowing which genes are involved in the biosynthesis of STLs or in the development of laticifers, and using CRISPR/Cas9 gene editing, it should be possible to rapidly breed novel varieties that have an altered STL profile, or with a reduced or increased laticifer volume. For example, inactivation of the genes encoding kauniolide synthase (KLS) resulted in lines accumulating free or conjugated costunolide, a potential anticancer STL (Cankar *et al.*, 2022). For chicory breeding efforts targeting laticifers or the production of metabolites in the laticifers, it is therefore essential to acquire basic information about the metabolite and transcriptome pattern of different plant tissues across developmental stages. However, chicory is not a model species and there are few transcriptomic, metabolomic and genomic data available until now (Testone *et al.*, 2017, Testone *et al.*, 2019, Peña-Espinoza *et al.*, 2020). Therefore, in this study we set out to determine the metabolite and transcription profiles of several tissues of chicory roots, including latex, which is the cytosolic compartment of the laticifers. The combined analysis of metabolome and transcriptome data has been very useful to determine key genes and regulatory processes in plants (Balcke *et al.*, 2017, Cao *et al.*, 2017a, Verhoeven *et al.*, 2017).

Analysis of the data reported here provides unique insights into the metabolic and development processes in chicory taproot. In particular, we could show that STLs are essentially stored in the latex but that their synthesis is compartmentalized across the root tissues and we identified candidate genes for the biosynthesis and transport of the chicory STLs as well as for the development of laticifers. This study therefore constitutes a foundation for further functional studies of these processes in chicory.

## Results

Morphological and cytological analysis showed the existence of different tissues and compartments across the taproots, including latex (LX), hypodermis (HD), cortex (CX), phloem (PH) and vascular cylinder (VC) (**Fig. S2**). It is difficult to properly dissect the CX and PH, but it is possible to dissect the HD and VC without contamination from the other tissues. We also collected the latex (LX), which is the cytosolic compartment of laticifer cells, by cutting plant roots and shoot parts. Therefore, in this study, we have used three different root tissues (LX, HD and VC) for targeted and untargeted metabolic profiling. In addition, we generated transcriptome data of LX, HD and VC to determine differentially expressed genes in these tissues from 16 week old plants.

### Metabolome profiling of STLs

#### STLs profiling of different root cell types

Although it is known that in latex-producing Asteraceae species, the STLs mostly accumulate in the latex (Sessa *et al.*, 2000, Seo *et al.*, 2009), there is no clear picture of the distribution of STLs in chicory roots. To analyse the pattern of STLs distribution across the plant tissues, we measured the peak area of STLs in three different plant tissues using a targeted approach (**Table S1**). The major STLs of chicory are lactucin (L), 8-deoxylactucin (DL), lactucopicrin (LC), lactucin-15-oxalate (LOx), 8-deoxylactucin 15-oxalate (DLOx) and lactucopicrin oxalate (LCOx) (**Fig. 1**).

**Figure 1.**
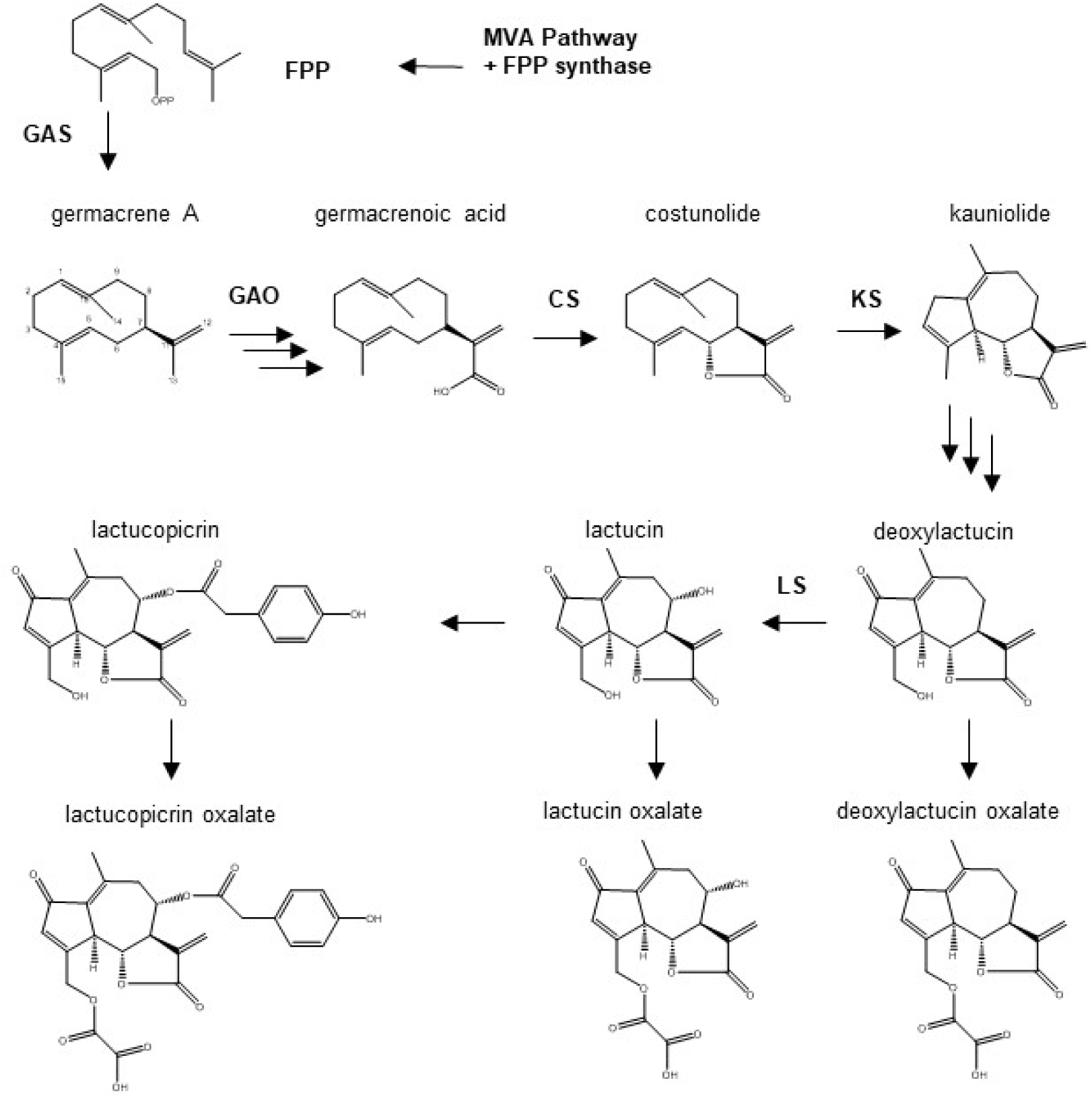
Biosynthesis pathway of sesquiterpene lactones in chicory. FPP: farnesyl diphosphate. The known enzymatic steps are abbreviated as follows: GAS: germacrene A synthase; GAO: germacrene A oxidase; COS: costunolide synthase; KLS: kauniolide synthase; LCS: lactucin synthase.

The STLs were measured from root sections by collecting material from three different tissues: LX, HD and VC (**Fig. S2**). In order to dissect different tissues from chicory root, it was necessary to distinguish them by staining of neighboring sections (**Fig. S2**). Principal component analysis of STLs in the three different root tissues shows a clear separation of the sample types (**Fig. 2A**), with STLs in latex being significantly higher than in other tissues (**Fig. 2B**). By contrast, the VC has the lowest concentrations of STLs. The oxalate forms of STLs are more abundant than non-oxalates in all cell types (**Fig. 2B**). Thus, our analysis of STLs in chicory roots confirms that they are largely stored in the latex, as shown in other related species.

**Figure 2.**
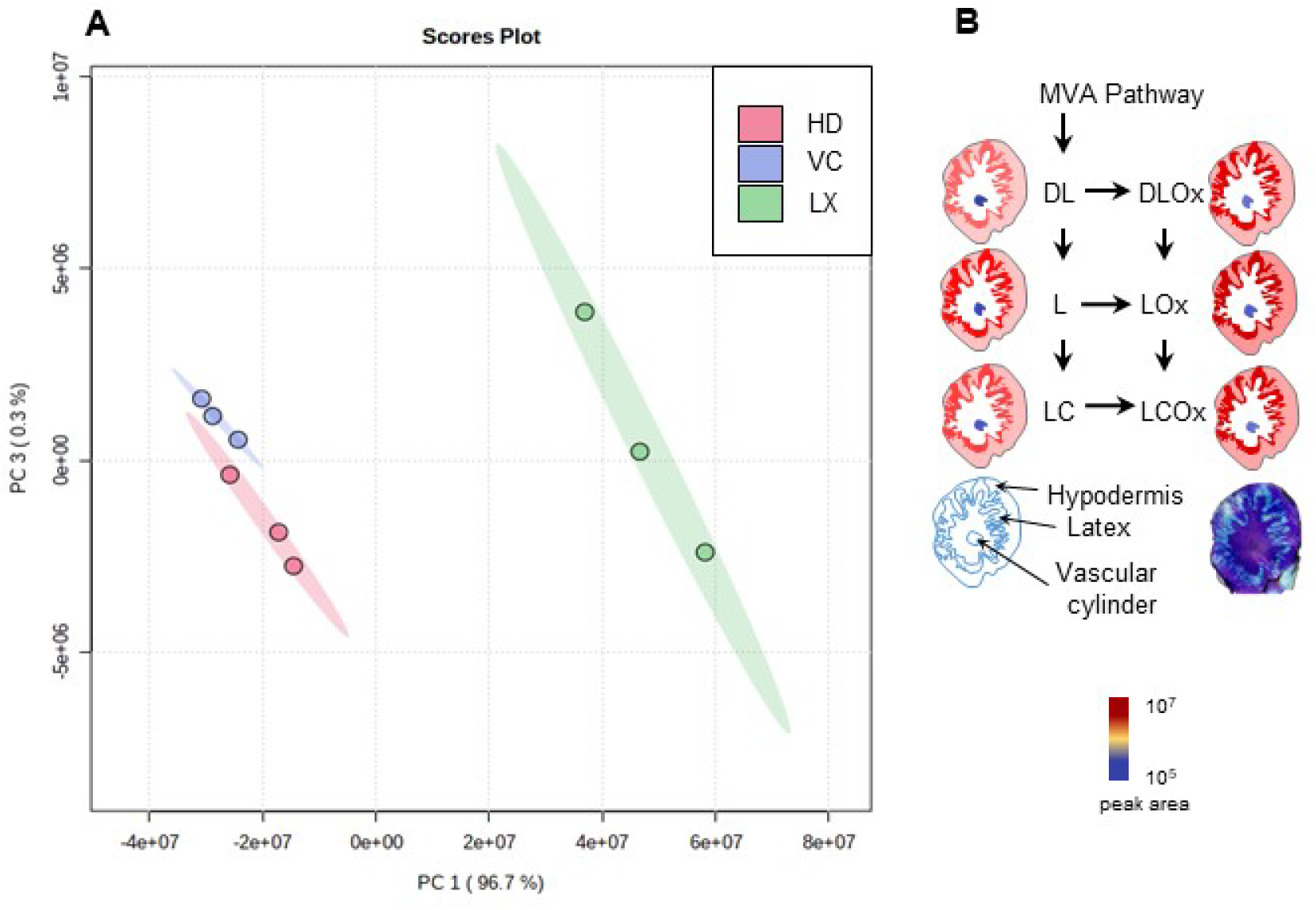
Profiling of sesquiterpene lactones (STLs) in chicory roots. **A.** Principal component analysis of biological repeats across different cell types based on the peak area of the major STLs using LC-MS positive mode (LC-MS-Pos). HD: hypodermis; VC: vascular cylinder; LX: latex. **B**. Distribution of the major STLs in different root tissues using targeted LC-MS positive mode. DL: deoxylactucin; DLOx: deoxylactucin-oxalate; L: lactucin; LOx: lactucin-oxalate; LC: lactucopicrin; LCOx: lactucopicrin-oxalate. The color scale indicates the peak area of the measured compounds normalized to tissue fresh weight.

### Untargeted metabolome profiling of chicory root tissues

To explore the metabolite distribution pattern across different cell types of chicory roots, we measured metabolites extracted from two tissues (HD and VC) and LX using exhaustive LC-MS and GC-MS profiling (**Fig. 3**). Untargeted LC-QToF-MS/MS was performed using both negative (LC-MS-Neg) and positive (LC-MS-Pos) ionization modes. In addition, we used an untargeted profiling approach for hydrophilic metabolites (LC-MS-hpm) (Balcke *et al.*, 2017). GC-MS was done with hexane extracts.

Principal component analysis (PCA) of the samples based on metabolome data using different MS approaches again showed clear separation of the samples according to tissue type, for which the LC-MS–neg dataset constitutes a representative example (**Fig. 3A**). A total number of 21 437 features were identified from the metabolome profiling data of chicory root tissues using five different methods. LC-MS-hpm with 12,954 and GC-MS with 1,657 features, produced the largest and smallest datasets, respectively. Annotation of the different datasets from various cell types identified a total number of 878 potential metabolites. The highest and lowest rates of annotation were for GC-MS and LC-MS-Pos, respectively. LC-MS-hpm producing the largest dataset (**Fig. S3**). In the GC-MS, LC-MS-hpm, LC-MS-Pos and LC-MS-Neg measurements, the most abundant compounds across all tissues are present in the LX samples. Based on the ClassyFire annotation (Djoumbou Feunang *et al.*, 2016), these compounds belong to lipids, fatty acids, steroids, sesquiterpene lactones and their derivatives. Also, across all tissues, the most abundant compounds in LC-MS-Neg measurements are present in the VC. Based on the ClassyFire annotation, these compounds mainly belong to the subclass of alcohols and polyols, amino acids and peptides, fatty acids and conjugates (**Fig. S3 and Table S2**).

In addition to STLs, other mevalonate pathway-derived metabolites such as squalene, triterpenes and sterols accumulated in the latex, similar to what was shown in Russian dandelion (*Taraxacum koksaghyz*) (Benninghaus *et al.*, 2020). Sterols and fatty acyls are involved in structural integrity and energy storage and occur in LX in association with rubber (Schulze *et al.*, 2011). These molecules are present in higher amounts in the LX of chicory roots compared to other tissues (**Fig. 3B** **and Table S2**).

**Figure 3.**
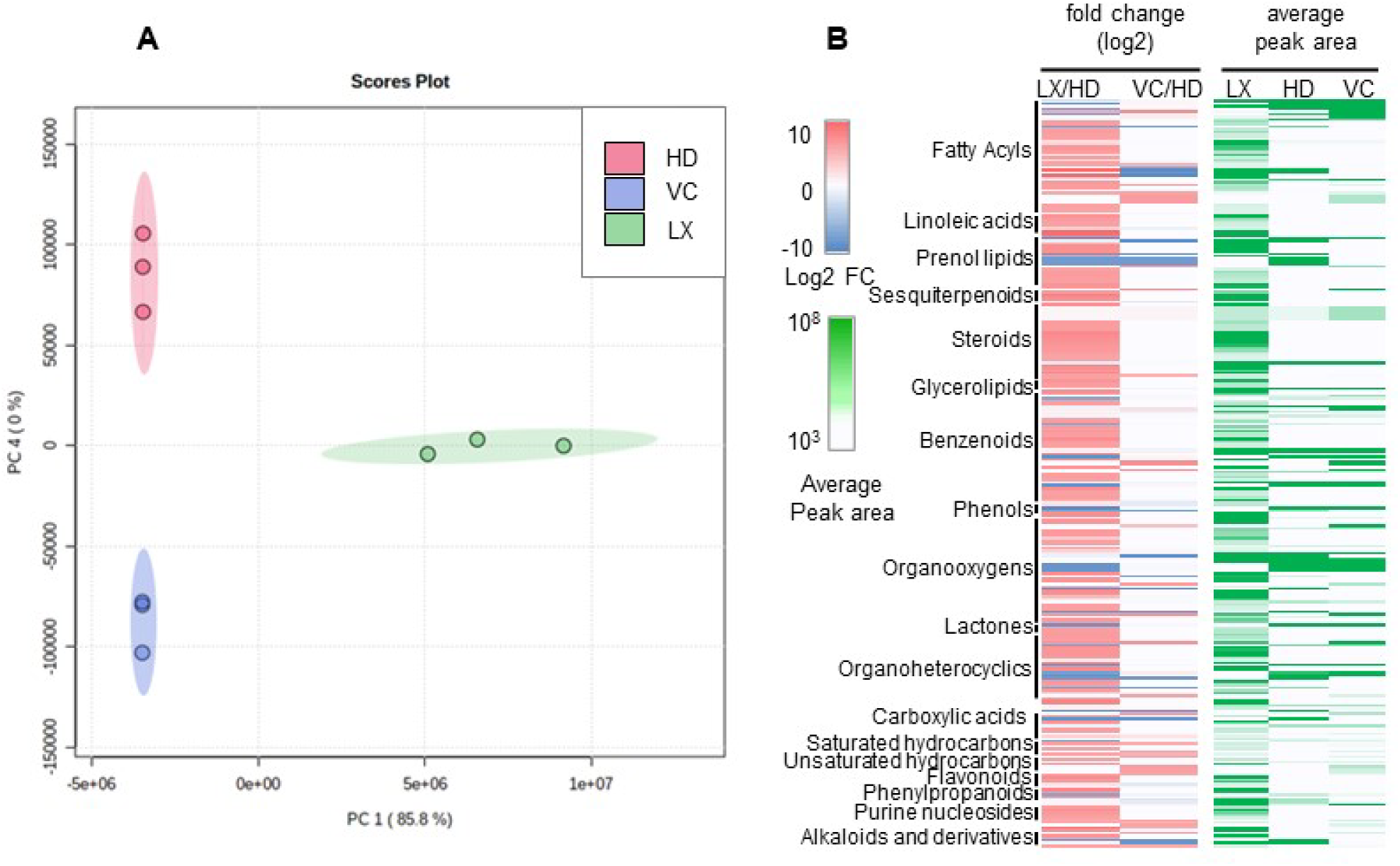
Metabolome profiling of chicory roots. **A.** Principal component analysis of various biological repeats across three root tissues based on the peak area from metabolite profiling using untargeted profiling (LC-MS-neg). **B**. Abundance (right side) and fold change relative to HD (left side) of classes of metabolites across three root tissues. LX: latex; HD: hypodermis; VC: vascular cylinder.

Pathway enrichment analysis of the annotated compounds indicates that the majority of the compounds map to the biosynthesis of secondary metabolites (n = 58). *cis*-1,4-polyisoprene is a major component of natural rubber and the rest consists of related proteins, carbohydrates, lipids, sterols and terpenoids (Schulze *et al.*, 2011). Consistent with this, pathways related to the biosynthesis of unsaturated fatty acids (19 features) and fatty acid biosynthesis (11 features) were enriched in LX. In total, the LX contains most of the compounds (75 features) mapped to metabolic pathways and VC holds the lowest numbers (9) (**Table S4**).

As shown in **Fig. S3**, the majority of the compounds could not be annotated using the MS-Dial or Xcalibur software programs. An enriched annotation of the metabolome is a necessary step for understanding it. Therefore, to associate the unknown compounds with known classes, we used MetFamily, a recently developed software that clusters MS features based on the similarity of their fragmentation patterns (Treutler *et al.*, 2016). For instance, we found, as expected, that the most abundant compounds detected in the latex by MS/MS in the negative mode belong to the group of STLs (**Fig. S4 and Table S4**).

### Transcriptome profiling

The percentage of mapped reads is an important mapping quality statistic since it is a global indicator of overall sequencing accuracy and the presence of contaminating DNA (Conesa *et al.*, 2016). For instance, depending on the read mapper utilized, 70-90 percent of normal RNA-seq reads will map onto the human genome (Dobin *et al.*, 2013). Despite availability of the genome for *Cichorium intybus*, Cultivar: Punajuju (ASM2352571v1), We decided to use de novo transcriptome assembly because of the lower mapping rate of O37 RNA sequencing reads on the genome of the Punajuju (<65%) in comparison to the de novo assembled transcriptome (>95%) (Figure S8) and also due to the lack of publicly available information about the Punajuju variety.

#### Tissue-specific gene expression based on RNA-Seq

We performed transcriptome analysis of the same root tissues used for the metabolome profiling, namely HD, VC and LX (**Fig. S2**). RNA samples were processed for RNA sequencing using Illumina sequencing. We performed differential gene expression analysis for the three tissues. The biological repeats of each of these tissues cluster tightly together and each cluster has a distinct expression pattern, supporting a reliable preparation of the tissues. LX has the highest number of tissue-specific genes (3784), followed by HD (1153) and VC (699). HD and VC have more commonly expressed genes (1736), followed by LX and HD (857) and LX and VC (447) (**Fig. 4A**). Mapping of annotated transcripts to pathways using KEGG mapper (Kanehisa, 2000) showed that the biosynthesis of secondary metabolites, hormone signal transduction, plant-pathogen interaction and phenylpropanoid biosynthesis were the most enriched processes across different cell types (**Table S4**).

**Figure 4.**
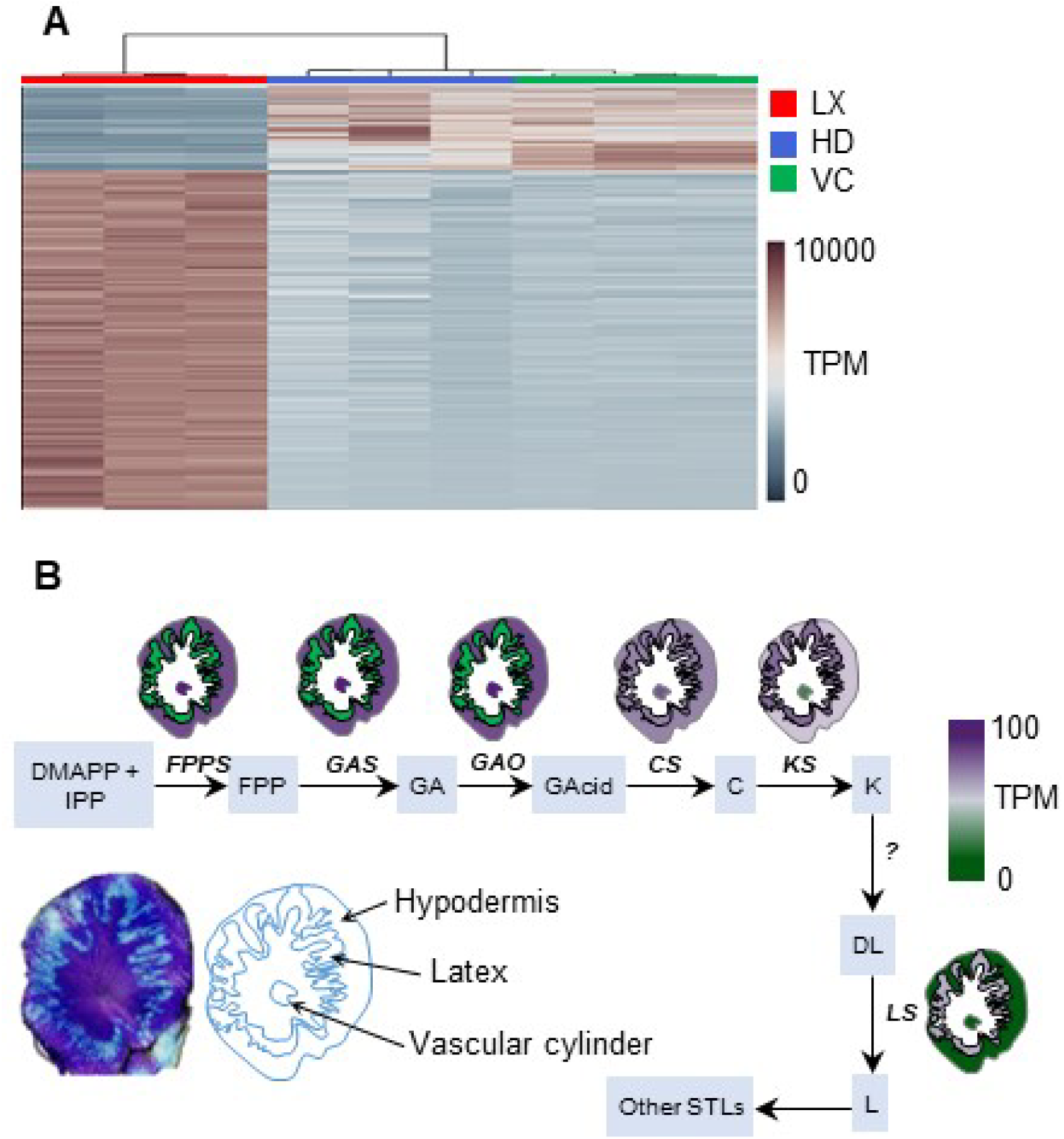
Transcriptome profiling of chicory root tissues. **A**. Clustering of transcriptome samples. **B**. Expression pattern of the known steps of STL biosynthesis across root tissues. C: costunolide, FPP: farnesyl diphosphate, GA: germacrene A, GAcid: germacrenoic acid, K: kauniolide, COS: costunolide synthase, DL: deoxylactucin, L: lactucin FPPS: farnesyl diphosphate synthase, GAS: germacrene A synthase, GAO: germacrene A oxidase, KLS: kauniolide synthase, LCS: lactucin synthase.

#### Biosynthesis of STLs occurs across several tissues

To gain further insight in the distribution of STL biosynthesis across root tissues, we collected gene expression data for the known genes of the pathway, namely farnesyl diphosphate synthase (*FPPS*), germacrene A synthase (*GAS*), germacrene A oxidase (*GAO*), costunolide synthase (*COS*) kauniolide synthase (*KLS*), and lactucin synthase (*LCS*). Our transcriptome data shows that genes involved in the early steps of STLs biosynthesis, including *FPPS*, *GAS* and *GAO,* are expressed in HD and VC. However, the next steps, *COS*, is expressed at similar levels in all tissues, while the downstream steps, (*KLS* and *LCS*), are preferentially expressed in the latex compared to HD and VC (**Fig. 4B**). This indicates that intermediates up to germacrenoic acid (GA) and costunolide (C) are synthesized outside the latex, and that germacrenoic acid and costunolide are then transported to the laticifer cells and there further converted to downstream products.

#### Differential expression of transporters across chicory root tissues

The compartmentalization of STL biosynthesis gene expression would necessitate the transport of STL pathway intermediates between root tissues. Indeed, we observe striking differential expression of distinct classes of ABC transporters. In the LX, ABCB/E/F are highly upregulated, while ABCC/G are mainly expressed outside of the LX (**Fig. 5, Table S5**). High expression of the ABCG/C outside of the LX might be an indication of their role in exporting metabolites and precursors into the LX. ABCG is the largest subfamily of ABC transporters, which mainly play roles in the export of metabolites (Wu *et al.*, 2014, Natarajan *et al.*, 2020, Banasiak *et al.*, 2021, Philippe *et al.*, 2022) such as cutin precursors, cutin monomers, wax, cuticular lipids, alkanes suberin (Xin and Herburger, 2021), lipid (Banasiak *et al.*, 2021), diterpenoids (Jasinski *et al.*, 2001) and toxic metabolites (Goldstone, 2008). Thus, these might be candidates for the export of GA and C from tissues peripheral to the LX. In contrast, ABCB transporters are involved in the import of metabolites such as auxin (Paterlini, 2020), secondary metabolites and other substances (Xie *et al.*, 2020). Thus, overexpression of ABCB transporters in the LX is consistent with an import of metabolites to the latex. This could be for example GA and C for the STL biosynthesis.

**Figure 5.**
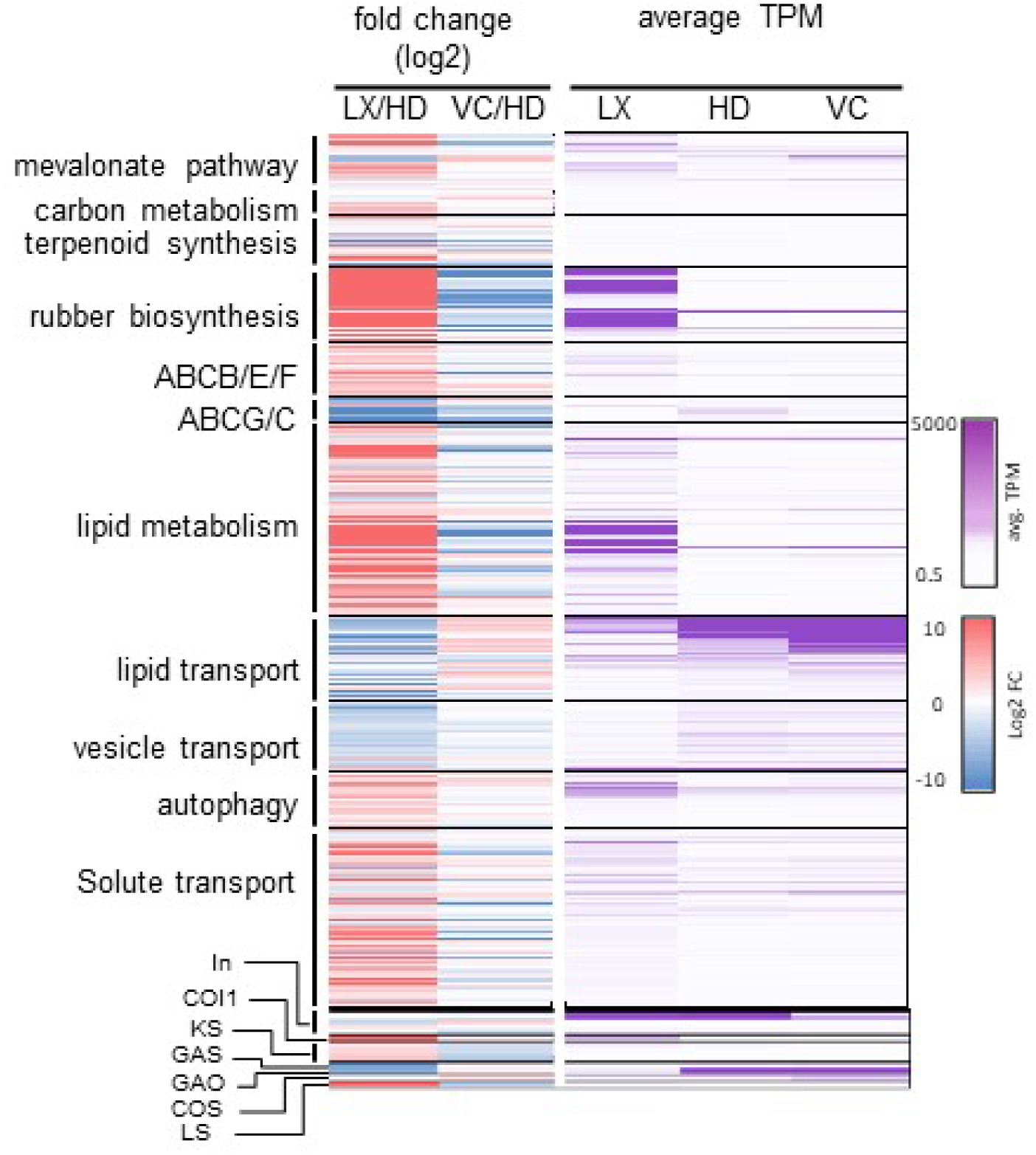
Expression data of selected pathways and gene families. Left panel: Log2 fold change of latex to hypodermis (LX/HD) and vascular cylinder to hypodermis (VC/HD). Right panel: gene expression values in transcripts per million (TPM). ABCB/E/F: ABC transporters of families B, E and F; ABCG/C: ABC transporters of families G and C; In: inulin biosynthesis; COI1: jasmonate receptor; KLS: kauniolide synthase; GAS: germacrene synthase; GAO: germacrene A oxidase; COS: costunolide synthase.

#### Genes involved in rubber and triterpenoid biosynthesis are overexpressed in latex

In Russian dandelion, which is related to chicory, in addition to STLs the latex contains triterpenoids and natural rubber, whose biosynthesis also requires isoprenyl precursors from the cytosolic mevalonate pathway. In rubber tree (*Hevea brasiliensis*) and Russian dandelion (*Taraxacum kok-saghyz*), natural rubber biosynthesis takes place in the latex (Cherian *et al.*, 2019). The biosynthesis of *cis*-1,4-polyisoprene takes place on the surface of rubber particles, with *cis*-polyprenyl transferases (CPTs) being the major enzymatic activity (Cherian *et al.*, 2019). A phylogenetic tree of CPT homologs present in chicory indicates that there are genes orthologous to known rubber CPTs from the related *Taraxacum* species (**Fig. S6**). CPTs alone cannot synthesize long chains of *cis*-1,4-polyisoprene but require the presence of additional proteins present in the rubber particles, including rubber small rubber particle proteins (SRPP). As expected, genes encoding CPTs and CPT-binding proteins (Epping *et al.*, 2015, Qu *et al.*, 2015) are specifically upregulated in the LX (**Fig. 5; Table S5**). Also, higher expression of squalene synthase and squalene epoxidase in the latex (**Table S5**) is consistent with the accumulation of triterpenoids in the latex, as is the case in the related *Taraxacum spp.* (Pütter *et al.*). Furthermore, genes encoding enzymes of the mevalonate pathway are strongly upregulated in the LX, consistent with its role in supplying precursors for the synthesis of natural rubber and triterpenoids (**Table S5**).

#### Gene expression for inulin metabolism and distribution of inulin and related sugars in chicory roots

The highest concentration of inulin is in HD and VC, whereas LX has the lowest content (**Fig. 6** **and Table S6**). Interestingly, sucrose, the building block of inulin, showed different distributions. Sucrose was most abundant in the vascular cylinder, with decreasing concentrations in the LX and HD. Glucose and fructose levels are relatively low compared to sucrose and especially low compared to inulin levels. Glucose is produced in the first step of inulin synthesis when SST catalyses the synthesis of the shortest inulin from two sucrose molecules. Fructose is produced when inulin is broken down by FEH activity. Glucose was more abundant in the HD, whereas fructose, surprisingly, was most abundant in the LX. Thus, fructose, which is a major ingredient of inulin, and released when inulin is degraded by FEH, is highest in latex which might point to breakdown of inulin in the LX. The genes for inulin biosynthesis, respectively encoding sucrose-sucrose 1-fructosyltransferase (SST) and fructan:fructan 1-fructosyltransferase (FFT), are most strongly expressed in the HD (**Fig. 6**). Nonetheless, these genes are also relatively high expressed in the LX and significantly less in the VC although inulin levels in VC are comparable to HD. This suggests that although expression of inulin biosynthesis genes seems to take place in LX, it might not lead to inulin biosynthesis enzyme activity in LX. (**Fig. 6**). However, the low levels of inulin in the LX combined with high levels of fructose suggest that inulin in the LX is rapidly degraded or exported to neighboring tissues. Interestingly, ABC transporters have also been shown to be responsible for the transport of inulin by bacteria (Tsujikawa *et al.*, 2021). Individual ABCG transporters that are overexpressed in LX (**Fig. 5**) constitute potential candidates for the export of inulin out of the laticifers. The expression of genes involved in the degradation of inulin encoding isoforms of fructan exohydrolase (FEH-I, FEH-IIa and FEH-IIb) is rather low but for FEH-IIa highest in LX and for FEH-IIa and b highest in HD, and most likely they play a role in the low inulin level found in latex.

**Figure 6.**
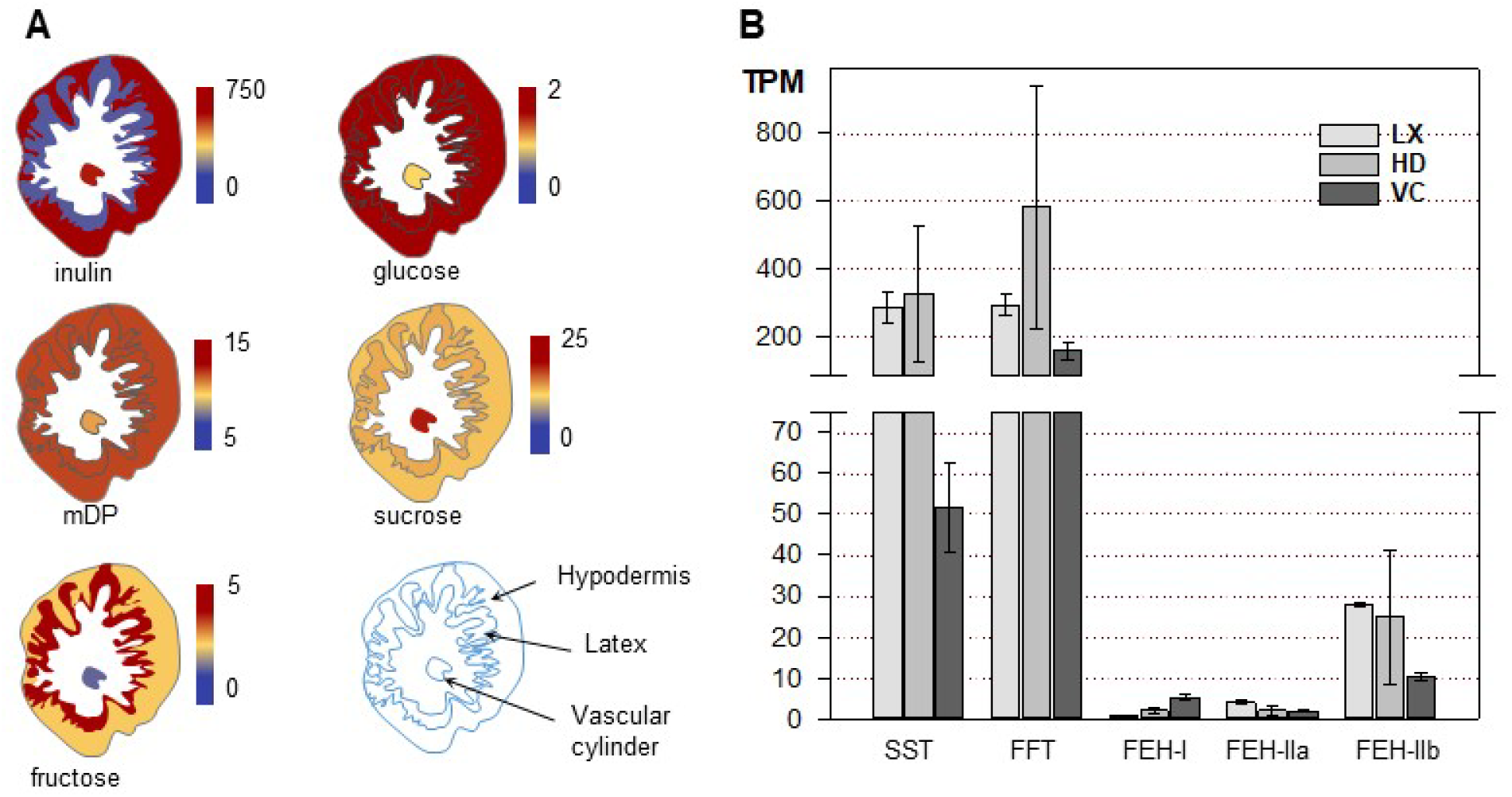
Inulin metabolism in chicory roots. A. Content of inulin and related sugars. mDP: mean degree of polymerization of inulin. Values are mg/g dry weight except for mDP. The data are available in Table S5. B. Inulin metabolism gene expression in chicory roots. SST: sucrose:sucrose 1-fructosyltransferase; FFT: fructan:fructan 1-fructosyltransferase; FEH: fructan exohydrolase. Average of transcripts per million (TPM) from three biological replicates ± standard error.

#### Hormone profiling shows increased jasmonates and abscisic acid levels in latex

There is little information available to date on phytohormone levels associated with laticifers in plants. We thus measured several phytohormones in LX, HD and VC with a focus on jasmonate (JA) related metabolites. Phytohormone levels for JA, OPDA, JA-isoleucine and abscisic acid (ABA) were significantly higher in LX relative to other tissues, while SA levels were not significantly different between tissues (**Fig. 7**). OPDA levels were particularly high in LX, reaching levels of several thousand ng/g FW. It has been known that Me-JA, OPDA and JA are involved in the modulation of the latex in the rubber trees (Hao, 2000, Duan *et al.*, 2010, Laosombut *et al.*, 2016). However, this was done either by supplying exogenous jasmonates or upon wounding of the plants (Hao, 2000).

**Figure 7.**
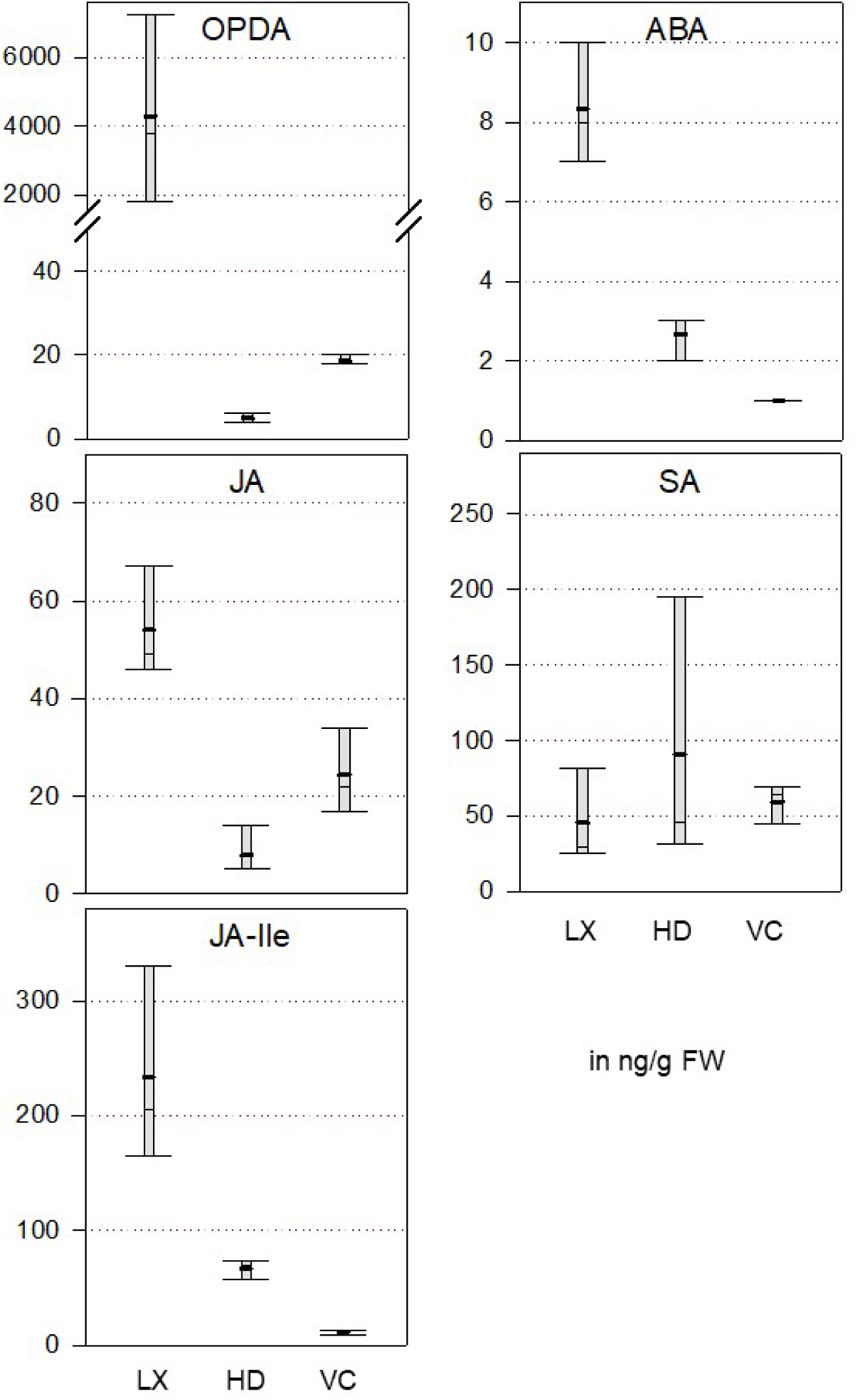
Concentration of different plant hormones in chicory root tissues. Concentrations are in ng/g fresh weight. JA: jasmonic acid; OPDA: 12-oxophytodienoic acid; JA-Ile: jasmonate isoleucine; ABA: abscisic acid; SA: salicylic acid. HD: hypodermis; LX: latex; VC: vascular cylinder. The data are based on measurements from three biological replicates. The box plots indicate the highest and lowest values, as well as the mean (thick black bar) and the median of the three measurements. Number of biological replicates = 3.

#### Other aspects

Analysis of our transcriptome data indicates the upregulation of several elements of ubiquitin-mediated protein degradation and protein degradation-related signaling pathways (**Table S4**). Protein turnover is necessary for tissues with high metabolic activity (Vierstra, 1993, Quigg and Beardall, 2003), which is the case in laticifers (Tungngoen *et al.*, 2009, Tang *et al.*, 2013). Moreover, up-regulation of aquaporins was also observed in LX. The aquaporin gene (HbPIP2;1, T11M8-2) was suggested to be involved in regulating the water transfer between the laticifers and surrounding tissues and its expression has been positively correlated with ethylene stimulation of latex yield (Tungngoen *et al.*, 2009). Also, it has been suggested that up-regulation of aquaporins is involved in the enhanced latex regeneration (Tungngoen *et al.*, 2009). Consistently, in our dataset, 20 aquaporin encoding transcripts have been upregulated in LX (**Table S5**).

#### Integrative analysis of transcriptome and metabolome data

Integrative analysis of metabolome and transcriptome data would provide a comprehensive picture of the regulatory pathways and uncover processes that are not obvious through analysis of the individual omics data (Sharma *et al.*, 2020). We thus performed co-analysis of the transcriptome and metabolome data by mapping them to the KEGG pathways (**Table S3 and Fig. S5**). Because the annotation of many metabolites is tentative and many metabolic features could not be annotated, this integrative analysis should be taken with caution.

Not surprisingly, metabolic pathways and the biosynthesis of secondary metabolites are strongly represented in LX. Noteworthy was the overrepresentation of vesicular transport and endocytosis in LX. This may be connected to the abundant metabolic pathways that occur in the laticifers, particularly rubber biosynthesis and may also reflect the fact that the laticifers constitute a sink tissue. Furthermore, the highest pathway enrichment in LX was for the synthesis and degradation of ketone bodies. Ketone bodies are degradation products of lipids, whose role in animals is to serve as a circulating energy source during calorie restriction (Cahill, 2006). They also have a signaling function in maintaining energy balance in animal cells (Newman and Verdin, 2014). In the context of chicory roots, ketone bodies could represent an alternative energy source when sugars (fructose and glucose) are limiting due to their use in inulin biosynthesis. Finally, the MAPK signaling pathway was also overrepresented in LX. Generally, MAPK signaling is involved in stress or hormonal responses (Taj *et al.*, 2010). Both could play a role in LX, either in the development or in the metabolic status of laticifer cells.

In the HD, we noted a strong enrichment in autophagy (**Table S4**). In Euphorbia, autophagy plays a role in the development of non-articulated laticifers through the fusion of autophagosomes and lysosome-like vesicles (Zhang *et al.*, 2018). Since laticifers emerge in the HD, this would also suggest a role for autophagy in the development of laticifers in chicory. The Glycosylphosphatidylinositol (GPI)-anchor biosynthesis (GPI-AP) pathway is one of the most enriched categories in both omics datasets in the HD (**Table S4**). GPI-AP is involved in facilitating trafficking routes and are anchored at the cell surface (Maeda and Kinoshita, 2011). This is also consistent with the high regulation of the ABCG/C in the HD (**Fig. 5**), which may be relevant for the transfer of metabolites to the LX. We note also the enrichment in several lipid biosynthesis pathways (sphingolipid, glycerolipid and unsaturated fatty acid) in HD. This again may have to do with the transfer of metabolites to the laticifer but also possibly with the storage of inulin in the vacuole (Darwen and John, 1989).

Finally, there is a strong enrichment of the categories fructose and mannose metabolism in all tissues (LX, HD and VC). This is most likely due to the biosynthesis of inulin, which is a major process in chicory roots.

## Discussion

In this work, we generated a comprehensive dataset of metabolomics and transcriptomics for different tissues of the chicory root. Chicory is a specialized crop for the high-scale production of inulin, which accumulates in large quantities in the taproot. In addition, chicory produces a milky latex that contains high amounts of sesquiterpene lactones (STLs) as well as *cis*-1,4-polyisoprene. The presence of the latex represents a problem for inulin processing but also a potential opportunity for the production of STLs, some of which have potential pharmacological applications (Peña-Espinoza *et al.*, 2015, Matos *et al.*, 2020, Peña-Espinoza *et al.*, 2020, Häkkinen *et al.*, 2021). Furthermore, Russian dandelion (*Taraxacum koksaghyz*) and guayule (*Parthenium argentatum*) produce a milky latex whose *cis*-1,4-polyisoprenehas properties that are similar to those of the rubber tree (*Hevea brasiliensis*) and constitute potential replacements for natural rubber but with a better allergenic profile. Thus, generating tissue-specific transcriptomics and metabolomics should provide insights both into the production of inulin but also on the development of the laticifers as well as the biosynthesis of compounds accumulating in the latex. Currently, such a comprehensive and well documented study for chicory and related species is not available. The most salient results of our multiomics dataset are discussed below.

### Biosynthesis of sesquiterpene lactones occurs across several tissues

STLs are the major bitter compounds associated with chicory taste and are one of the reasons for using chicory as a coffee replacement. The most abundant STLs accumulate in the LX, particularly the oxalate derivatives of deoxylactucin, lactucin and lactucopicrin, which collectively represent the end-products of the pathway. Interestingly, intermediates such as costunolide, for example, could not be detected, indicating a complete conversion to the late products. Only in lines where genes encoding kauniolide synthase (KLS) were knocked-out could costunolide (C) be detected, but mostly cysteine or glutathione conjugates of C (Cankar *et al.*, 2022). Our gene expression data indicate that up to germacrenoic acid (GA) and C, the biosynthesis takes place outside the LX and that from C onwards, the biosynthesis is localized in the latex. These data are consistent with those of previous studies, where the expression of genes for GA synthase (GAS), GA-oxidase and costunolide synthase (COS) investigated by RT-qPCR was shown to be low in the latex (Kwon *et al.*, 2022) and promoter:reporter gene fusions confirmed the expression of GAS in the root cortex (Bogdanović *et al.*, 2019, Kwon *et al.*, 2022). The distribution of pathways for specialized metabolites across several tissues is not uncommon in plants. For example, in the early enzymatic steps of the opium poppy (*Papaver somniferum*), benzylisoquinoline alkaloids that accumulate in the latex occur in the sieve elements of the phloem and the corresponding genes are transcribed in the companion cells (Onoyovwe *et al.*, 2013, Ozber and Facchini, 2022). Similarly to the situation in chicory, the late steps of benzylisoquinoline alkaloid biosynthesis do occur in the latex (Onoyovwe *et al.*, 2013, Chen *et al.*, 2018). Additionally, some steps in benzylisoquinoline alkaloids biosynthesis do occur in both tissues, indicating that several intermediates may be transported from sieve elements to the latex (Onoyovwe *et al.*, 2013). In chicory, our knowledge of the STL pathway is still incomplete. So far, after germacrenoic acid, three steps are known: COS, KLS (Cankar *et al.*, 2022) and lactucin synthase (LCS) (Cankar *et al.*, submitted). COS is expressed at similar levels in all tissues, whereas KLS and LCS are specifically expressed in the latex. Still missing are oxidases for positions 2 and 15, a transferase for the conjugation of oxalate at position 15 and a transferase for the transfer of hydroxyphenyl acetate on position 8 for lactucopicrin. The oxidases are likely cytochrome P450 oxygenases of the CYP71 family, as is the case for most oxidases identified so far in STL biosynthesis from the Asteraceae (Teoh *et al.*, 2006, Liu *et al.*, 2014, Frey *et al.*, 2018, Liu *et al.*, 2018, Frey *et al.*, 2019, Frey *et al.*, 2020). A phylogenetic tree of CYP-encoding genes differentially expressed in latex (**Fig. S7**) reveals that indeed most belong to the CYP71 family and are therefore likely candidates for the missing steps in the pathway. If confirmed, this would suggest that most steps after costunolide are indeed localized in the latex. Thus, although transport of other intermediates cannot be excluded, it is probable that GA and C are the major intermediates transferred from surrounding tissues to the latificer cells.

### Implications for transport

In opium poppy there is indication that the transfer of intermediates from sieve elements to laticifers is apoplastic rather than symplastic. The recent identification of a small group of purine-permeases involved in the import of benzylisoquinoline alkaloids into the laticifer cells confirms this hypothesis (Dastmalchi *et al.*, 2019). Whether such permeases are also involved in the import of GA or C into laticifers is questionable, however, because sesquiterpenoids are structurally very different from alkaloids. Instead, we noted differential expression for a number of ABC transporters. Fittingly, a number of ABCG transporters, which are known to be exporters, are overexpressed in the HD, while ABCB transporters, which are in general importers, are overexpressed in the LX. The latter could therefore be involved in the import of GA or C into the laticifers, while the former would export GA or C from the HD. Validation of the function of these transporters could be done by creating knock-out lines by gene editing. However, the redundancy within the ABC transporter family, as visible from the number of transporters differentially expressed in chicory, may require inactivating several genes to see a noticeable effect.

### Implications of the distribution of inulin and sugars in the root tissues

Analysis of inulin content across the root tissues clearly shows that it accumulates throughout the root, except in the LX, (**Fig. 5**). Therefore the substrates for inulin biosynthesis, namely sucrose, which is imported from the leaf, would have to be transported from the phloem across the laticifers. The high concentration of fructose in the LX and the relatively high expression of inulin degradation genes in the LX suggest that some inulin is degraded in the LX, but that both inulin and fructose might also be exported to neighboring tissues. However, the degree of polymerization of inulin in the LX is similar to that of the HD (**Fig. 5**). This suggests that inulin is not extensively broken down in the LX but exported, or that shorter chain inulins are exported. One way to clarify this issue would be by identifying the trafficking mechanisms of inulin between tissues. In plants these are not known, but bacterial ABC transporters were shown to be involved in the import of inulin (Tsujikawa *et al.*, 2021), indicating that again ABC transporters may also be involved in the trafficking of inulin. Further studies will be needed to investigate the dynamics of inulin biosynthesis and transport in chicory roots during development.

### Composition of the latex

The latex is a complex mixture of multiple compounds. As mentioned above, it contains small molecules in quite large amounts, such as sesquiterpene lactones and triterpenoids. However, a typical component of latex is rubber (poly-*cis*-1,4-isoprene), which plays an important role in sealing wounds via coagulation mediated by oxidases (Wahler *et al.*, 2009). To the best of our knowledge, there is no report available on the presence and quality of rubber in chicory. As mentioned in the introduction, related Asteraceae species like Russian dandelion, guayule or lettuce have been proposed as alternative sources for natural rubber instead of the rubber tree, therefore, it is likely that chicory also produces rubber. This is confirmed by the strong overexpression in the LX of genes typically associated with rubber production, namely *cis*-polyprenyl transferase (CPT), rubber elongation factor and small rubber particle protein (**Fig. 5; Table S4)**. The length and amount of rubber produced by chicory are not known, however, and it would be worth investigating this further to evaluate the potential of chicory as a rubber producer. In contrast to Russian dandelion, the cultivation of chicory is well established in the north of Europe. It is also a highly productive crop with large taproots, although it has been bred for the production of inulin. Increased latex production could also be a breeding target, which may be more accessible nowadays with the availability of genome editing technologies. This will require deeper insight into the molecular processes that regulate laticifer development and the biosynthesis of rubber. One aspect of this is the hormonal regulation of development and biosynthetic processes.

### The role of phytohormones in laticifer formation and latex production

In *H. brasiliensis,* wounding and application of JA, methyl jasmonate (MeJA), linolenic acid and coronatine result in stimulation of laticifer formation (Hao, 2000, Tian *et al.*, 2015, Zhang *et al.*, 2016, Loh *et al.*, 2019). It has been suggested that JA promotes the differentiation of laticifer cells (Hao, 2000, Tan *et al.*, 2014) via the enrolment of JAZ, NINJA, TOPLESS (TPL), 28S proteasome, *SKP1*, *MYC1*, *MYC2*, and *EREBP1* in mature rubber trees (Chen *et al.*, 2011, Yang *et al.*, 2011, Zhao *et al.*, 2011). Also, it has been shown that expression of *COI1*, *MYB*, *ARF8* and *HB8* is correlated with MeJA-mediated differentiation of the laticifers (Laosombut *et al.*, 2016). However, all these data are based on artificial exposure to jasmonates, either by wounding or by applying exogenous JA or related substances to rubber trees. Thus, it is still unknown whether jasmonate signaling is involved in the developmentally programmed development and differentiation of laticifers. Several pieces of data in this work point to such a role for jasmonates in chicory. First, the levels of JA, OPDA and, importantly, JA-Ile are strongly elevated in the LX compared to other tissues (**Fig. 7**). OPDA is several hundred fold higher in LX than in HD or VC and JA-Ile is 3.5 and 20-fold higher in LX than in HD and VC, respectively. Additionally, in our transcriptomics data, several JA-related genes are upregulated in the LX (**Table S4)**. Most remarkably, we see a strong upregulation of a gene encoding a homolog of coronatine-insensitive protein 1 (COI1), which is the JA-Ile receptor and several transcription factors (TF) including a NAC, several DOF, a MYC and a WRKY TF (**Table S5**). In the rubber tree, *hbNAC1* is induced by wounding and dehydration and shown to activate the expression of an *SRPP* gene (Cao *et al.*, 2017b).

In the latex, the majority of JAZ-encoding genes have been slightly upregulated or show no regulation. Only JAZ8 is downregulated in the LX (**Table S4**). Unlike other JAZs, JAZ8 does not possess a canonical degron (Shyu *et al.*, 2012). The JAZ degron is a conserved LPIAR motif that blocks the binding site of JA-Ile and prevents the formation of the COI1-JAZ complex (Sheard *et al.*, 2010). Consequently, JAZ8 suppresses JA-regulated responses when expressed ectopically in *Arabidopsis* (Shyu *et al.*, 2012). Therefore, it is possible that the chicory JAZ8, which is down-regulated specifically in the latex, plays a role as a negative regulator in the process of laticifer development and differentiation.

## Conclusion

This study provides new insights into the metabolism and gene expression of chicory roots and reveals a complex interplay of metabolic and developmental processes. From this dataset, it is possible to predict candidate genes involved in the biosynthesis of latex components (STLs, rubber and triterpenoids) and in their transport and regulation, in laticifer development, and inulin metabolism and transport. Because chicory does not benefit from extensive genetics, in particular due to its being self-incompatible, the candidate genes identified by omics approaches can be tested with novel genome editing techniques, thereby bypassing the need for lengthy breeding approaches. This should lead to rapid improvements of the chicory crop, not only for inulin production but also for the development of new products based on the latex content.

## Materials and Methods

### Plant materials

A clone from *Cichorium intybus* (AA, 2n = 18, root chicory), Orchies 37 (O37) (Cankar *et al.*, 2021), was used and propagated via tissue culture and used in all the experiments described here. For the analysis of the STLs in plant tissues, 16-week-old plants grown at 25 °C, 16-h photoperiod, and light intensity of 80 mol/s/m2 was used (**Figure S2**).

### Microscopy of the laticifers and dissection of the cell types

We found that staining with toluidine blue O (TBO) differentiates various cell types. Hand sections of chicory roots (3 mm thickness) were stained with TBO (0.2 % in water) for 5 minutes and washed with distilled water. These stained sections were used as a guide to facilitate the dissection of different cell types in non-stained sections. Unstained subsequent sections were used for dissecting the HD and VC. LX was collected from roots.

### Targeted profiling of the STLs

Different tissue types (HD and VC) and latex from three different biological repetitions, the samples were frozen in liquid nitrogen. The tissue material (VC and HD) were ground in a mortar and all samples including latex were stored at -80°C. For sample preparation, 100 mg of each sample was transferred to a cryo-tube (for details see Balcke *et al.*, 2017), and the material was homogenized in a FastPrep instrument using an extraction buffer (80% MeOH, 19.9% H_2_O, 0.1% formic acid) and kept on ice. STLs identification by multiple reaction monitoring (MRM) was carried out by LC-MS/MS analysis of 5 µL of methanolic extracts using an Acquity UPLC (Waters) and a Sciex 6500 QTRAP and the software Analyst 1.7 (Sciex). Peak area quantification of six different STLs was done by the software Multiquant 3.0 (Sciex) The data and MRM parameters are provided in **Table S1**.

### Untargeted metabolome profiling

For untargeted metabolite analysis, a modified 2-phase extraction was performed where 900 µL dichloromethane/ethanol (2:1, -80°C) were first added followed by 150 µL dilute HCl (pH 1.7) was added to 100 mg of different tissues and extracted using the FastPrep instrument. Phase separation was done by centrifugation (2 min, 10,000 x g), the upper phase (containing hydrophilic metabolites) was collected and stored on ice. Then another 100 µL aqueous HCl was added, and the extraction and phase separation were repeated. Both aqueous extracts were combined and stored at -80 °C until untargeted analysis of hydrophilic compounds (see below).

The remaining organic phase was collected and stored on ice, and the tissue debris, was re-extracted with 500 µL tetrahydrofuran (THF). Both organic extracts were combined, dried in a nitrogen stream and dissolved in 80% methanol, centrifuged and analyzed according to Separation of medium polar metabolites (see below).

<u>Separation of hydrophilic metabolites</u> (hpm) was performed on a Nucleoshell RP18 (2.1 x 150 mm, particle size 2.1 µm, Macherey & Nagel, GmbH, Düren, Germany) using a Waters ACQUITY UPLC System, equipped with an ACQUITY Binary Solvent Manager and ACQUITY Sample Manager (10 µL sample loop, partial loop injection mode, 5 µL injection volume, Waters GmbH Eschborn, Germany). Eluents were A (aqueous 10 mmol/L tributyl amine, adjusted to pH 6.2 with glacial acetic acid) and B (acetonitrile), respectively. Elution was performed isocratically for 2 min at 2% eluent B, from 2 to 18 min with linear gradients to 36% eluent B, from 18-21 min to 95% eluent B and isocratically from 21 to 22.5 min at 95% eluent B, from 22.51 to 26 min at 2% eluent B. The flow rate was set to 400 µL/min and the column temperature was maintained at 40 °C. Metabolites were detected by negative electrospray ionization and mass spectrometry. The MS detection was done on a QTOF instrument in the negative mode (LC-MS-hpm) (see below).

<u>Separation of medium polar metabolites</u> (mpm) was performed on a Nucleoshell RP18 (2.1 x 150 mm, particle size 2.1 µm, Macherey & Nagel, GmbH, Düren, Germany) using a Waters ACQUITY UPLC System, equipped with an ACQUITY Binary Solvent Manager and ACQUITY Sample Manager (20 µL sample loop, partial loop injection mode, 5 µL injection volume, Waters GmbH Eschborn, Germany). Eluents A and B were aqueous solutions of 0.3 mmol/L NH4HCOO (adjusted to pH 3.5 with formic acid) and acetonitrile, respectively. Elution was performed isocratically for 2 min at 5% eluent B, from 2 to 19 min with a linear gradient to 95% B, from 19-21 min isocratically at 95% B, and from 21.01 min to 24 min at 5% B. The flow rate was adjusted to 400 µL/min and the column temperature was maintained at 40 °C. Metabolites were detected by positive and negative electrospray ionization using an Acquity UPLC (Waters) and TripleTOF 5600 mass spectrometer. The MS detection was done on a QTOF instrument in the negative mode and positive modes (LC-MS-neg and LC-MS-pos) (see below).

<u>Mass spectrometric analysis</u> of small molecules was performed by two strategies: targeted MS/MS via multiple reaction monitoring (QTRAP6500) and untargeted via MS-TOF-SWATH-MS/MS (TripleToF 5600, both AB Sciex GmbH, Darmstadt, Germany) operating in negative or positive ion mode and controlled by Analyst 1.7.1 and 1.6 TF software (AB Sciex GmbH, Darmstadt, Germany). The source operation parameters were as follows: ion spray voltage, - 4500 V/+5500 V; nebulizing gas, 60 psi; source temperature, 450 °C (QTRAP) or 600 °C (TripleToF); drying gas, 70 psi; curtain gas, 35 psi. For APCI, a nebulizer current of 3 units was used. TripleToF instrument tuning and internal mass calibration were performed every 5 samples with the calibrant delivery system applying APCI negative or positive tuning solutions, respectively (AB Sciex GmbH, Darmstadt, Germany).

Triple-ToF data acquisition was performed in MS^1^-ToF mode and MS^2^-SWATH mode. For MS^1^ measurements, ToF masses were scanned between 65 and 1250 daltons with an accumulation time of 50 ms and a collision energy of 10V (-10V). The MS^2^-SWATH-experiments were divided into 26 Dalton segments of 20 ms accumulation time. Together, the SWATH experiments covered the entire mass range from 65 to 1250 daltons in 48 separate scan experiments, which allowed a cycle time of 1.1 s. Throughout all MS/MS scans, a declustering potential of 35 (or -35 V) was applied. Collision energies for all SWATH-MS/MS were set to 35 V (-35) and a collision energy spread of ±25V, maximum sensitivity scanning, and otherwise default settings.

### Metabolome annotation

MS-Dial (v. 4.9) was used to extract and align MS1 and MS/MS features from SWATH-QToF runs (Tsugawa *et al.*, 2015). For best match spectra annotation of individual mz;r.t.-features an in-house MS/MS database was used in the MS-Dial environment. Xcalibur software was used to annotate GC-MS data using the NIST17 EI-MS library. The MetFamily (Treutler *et al.*, 2016) software program was used to annotate groups of features having similar MS/MS spectra in the LC-QToF-MS/MS data sets. With unique structural identifiers, such as SMILES supplied by database matching we generated ChemOnt terms of metabolite families based on the ClassyFire annotation (Djoumbou Feunang *et al.*, 2016). Metabolite functional classification was carried out using the KEGG Mapper (Kanehisa and Sato, 2019) online tool and enrichment Fold changes based on KEGG pathways were carried out according to the ratio of the proportions of the treatment to the background (Huang *et al.*, 2008).

### GC-MS analysis

100 mg of LX, HD and VC tissues were mixed with 2 ml hexane and after 5 minutes of centrifugation at 10,000 RPM, the hexane extracts were sealed in the GC vials. 1µl of extracts were injected directly into the GC–MS for analysis on a Trace GC Ultra gas chromatograph coupled to an ATAS Optic 3 injector and an ISQ single quadrupole mass spectrometer (Thermo Fisher Scientific) with electron impact ionization. The injection temperature rose from 60 °C to 250 °C at 10 °C/s and the flow rate of helium was 1 mL/min. The GC oven temperature ramp was as follows: 50 °C for 1 min, 50 °C to 300 °C with 7 °C/min, 300 to 330°C with 20 °C/min, and 330 °C for 5 min. Mass spectrometry was performed at 70 eV in full scan mode with *m/z* ranging from 50 to 450. Data analysis and chromatograms were implemented with the device-specific software Xcalibur (Thermo Scientific).

### Inulin and small sugar analysis

Carbohydrate analysis was performed to determine the concentrations of free fructose, sucrose, glucose, the inulin content and inulin mean degree of polymerization (mDP). Free sugars were extracted from 30mg freeze-dried root material in 800 µL Phosphate buffer (20mM, pH 7) at 85°C for 30 minutes using intermittent agitation. After centrifugation for 15 minutes at 14.000 rpm, the supernatant was collected and the pellet was resuspended in 400 µL Phosphate buffer and the extraction procedure was repeated. Supernatants were pooled and samples were de-ionised with a mixture of Q-Sepharose and S-Sepharose beads in Phosphate buffer (20mM, pH7). Samples were incubated for 5 minutes while shaking. After centrifugation, supernatant was collected and analysed. Free glucose, fructose, and sucrose concentrations were determined by ion chromatography using a Thermo Fischer ICS 5000 + DC system (Thermo Fischer Scientific, Waltham. USA) equipped with a Carbo Pac PA-1 column (4-250 mm) and a 25 μL sample loop, as described by (Stoop *et al.*, 2007) with some modifications. Monosaccharides were eluted from the column with a flow rate of 1 mL/min using a NaOH gradient: from 0–10 min, the concentration of NaOH in the eluent increased from 20 mM to 50mM in a linear manner; from 10-20 min a linear gradient from 50 mM to 330 mM was applied, followed by an isocratic elution using 330 mM NaOH for 5 minutes. An ED40 electrochemical detector fitted with a pulsed amperometric cell was used. Quantification of the three sugars was achieved by comparison of peak areas to external glucose, fructose and sucrose standards (Merck KGaA, Darmstadt, Germany). The inulin content was analysed in hydrolysed extracts. Hydrolysis was performed by incubating 30 µL de-ionised sample with 30 µL of 200 mM HCl for 2 hours at 60°C. Glucose and fructose concentrations were determined by ion chromatography as described for free sugars. The inulin content was calculated as the total carbohydrate content from which the concentrations of free glucose, fructose, and sucrose were subtracted. The mDP was calculated as the sum of bound glucose and fructose, divided by de amount of bound glucose.

### RNA isolation and sequencing

O37 lines root tissue at the 16-week age used for dissection of the different cell types from root sections. Dissected tissue and harvested latex from 3 independent biological repeats were supercooled in liquid nitrogen. Different cell types were grinded to a fine powder in presence of liquid nitrogen and transferred to -80 C frizzier. 100 mg of biomass was used to isolate total RNA by RNAeasy kit (QIAGEN) NanoDrop and BioAnalyzer 2100 (Agilent Technologies Inc) used for evaluation of RNA concentration and RNA integrity number (RIN > 7) subsequently.

RNA was sequenced in 150 bp paired-end mode using an Illumina NovoSeq 6000 (Novogene).

RNAseq data sets have been listed in the NCBI (www.ncbi.nlm.nih.gov) as a BioProject with accession number of PRJNA824299.

The gene expression pattern of different tissues from chicory root was elucidated in various types of root cells, including latex, HD and VC. Dissection of different tissues from chicory root carried out with the guides of stained sections (**Figure S2**). For this aim, paired sections with a diameter of 2 mm were selected; one was stained by TBO to differentiate various cell types and other was used for dissection. The NGS transcriptional analyses of different cell types were produced via RNA sequencing of the same material that was used for the LC-MS study (this provides the possibility of doing transcriptome and metabolome data co-analysis). The five different tissues (100 mg) were dissected and latex was collected as explained above from sixteen-week old plants. Three biological replicates for each tissue (HD, VC) and latex (in total 9 samples) were generated for RNA extraction. Collected biomass was used to isolate total RNA using the RNAeasy kit (QIAGEN). The RNA concentration (estimated by Nanophotometer NP80, Implen) and RNA integrity number (RIN > 8) were determined (by BioAnalyzer 2100, Agilent Technologies Inc.). RNA was sequenced (from nine RNA samples) in 100 bp strand-specific mode using an Illumina sequencing system. Samples were sent for sequencing in October 2018.

### Transcriptome assembly, functional annotation and gene expression analyses

The quality of raw reads was examined by FastQC v 0.11.5 (https://www.bioinformatics.babraham.ac.uk/projects/fastqc/). Trimmomatic v0.32 (Bolger *et al.*, 2014) was used for cleaning up the paired-end Illumina raw reads from low-quality and sequencing adapters. *de novo* approach using Trinity (Grabherr *et al.*, 2011) used for assembly of the high-quality reads (default parameters). bowtie2 (Langmead and Salzberg, 2012) and RSEM (Li and Dewey, 2011) were used to map the trimmed reads on the Trinity assembly and estimate the abundance of the read count subsequently. Expression values were normalized and measured as TPM (transcript sequence per million of mapped reads) values.

Bowtie2 was used to map the filtered reads back to assembly. RSEM was used to estimate the transcript abundance. All statistical analysis is carried out using EdgeR. We used quantitative PCR to confirm the results of RNA-seq data for all interested genes.

Paired-end Illumina raw reads were filtered by cutadapt and Trimmomatic v0.32 (Bolger *et al.*, 2014) with quality trimming to eliminate adapter contamination and low-quality reads. Fastq and MultiQC were used to evaluate the quality of the reads before and after trimming. Assembly was carried out using Trinity (Grabherr *et al.*, 2011).

Transcriptome annotation was carried out using in-house BlastX (cut-off *E*-value of 10–5) against the following databases: Nr, NCBI non-redundant database (January 12, 2015); TAIR, The Arabidopsis Information Resource (TAIR10); SwissProt, TrEMBL, UniProt Knowledgebase (UniProtKB) and KOG (eukaryotic orthologue groups) (Koonin *et al.*, 2004). Gene ontology and KEGG (Kanehisa, 2000) annotations were generated by Blast2GO (Conesa *et al.*, 2005) using BLASTx hits mining the Nr database. GO functional classification was carried out using DAVID (Huang *et al.*, 2008) KEGG Mapper (Kanehisa and Sato, 2019) online tools.

### Statistical analyses

The principal component analysis (PCA) was applied to samples of different cell types normalized to fresh weight in metabolome and transcriptome studies to evaluate the distribution and grouping among samples. CA plotted in R 3.6.1 using mean-centered and standardized data (unit variance scaled). Differential expression analysis was carried out by the edgeR package (Robinson *et al.*, 2009).

A trimmed mean of M values (TMM) normalization was applied to all samples during transcriptome data analysis. A threshold of log_2_ fold change ratio (log_2_ FC) ≥ 1 was used to define significant gene expression differences after cutting off the data at false discovery rate (FDR) value ≤ 0.05 for all unigenes with more than or equal to 1 TPM.

Gene ontology (GO) terms and Kyoto Encyclopedia of Genes and Genomes (KEGG; www.genome.jp/kegg/pathway.html) pathway analysis were carried out to identify biological functions and pathway involvement of DEGs with statistically significant differences; at P<0.05 was considered to indicate a statistically significant difference (Kanehisa, 2000). P ≤ 0.05. The fold enrichment (FE) was calculated based on Huang *et al.* (2009) as the ratio of the proportions of the treatment to the background.

Co-analysis of the metabolome and transcriptome data was carried out using submitting respected annotated features to color tool in the KEGG Mapper (Kanehisa and Sato, 2019) Fold changes based on KEGG pathways were calculated according to Huang *et al.*(2009) as the ratio of the proportions of the treatment to the background. Since the pathway mapping files of *Cichorium intybus* have not been introduced in the KEGG, we have used *Lactuca sativa* for the mapping of the data to the pathways.

## Data availability statement

The transcriptome data presented in this study are deposited in the NCBI repository under accession numbers PRJNA824299.

The metabolome data are deposited in the Metabolights database under the accession number MTBLS6917 (www.ebi.ac.uk/metabolights/MTBLS6917)

## Acknowledgments

This work was funded by the European Commission, Horizon 2020 Programme, Research and Innovation Programme, under grant agreement No. 760891 (H2020-NMBP-BIOTEC-07-2017: New plant breeding techniques (NPBT) in molecular farming: multipurpose crops for industrial bioproducts).

## Author contributions

AT and KV designed the experiments. KV produced and analyzed the transcriptome and metabolome data. GUB provided supervision for the production and analysis of the metabolome data. BA processed and analyzed the transcriptome data. JCH and IVdM analyzed the inulin and sugar content. AT and KB wrote the manuscript, which was proofread and approved by all authors.

