## Supplemental Figures for "Metabolome and transcriptome profiling of root chicory provide insights into laticifer development and specialized metabolism"

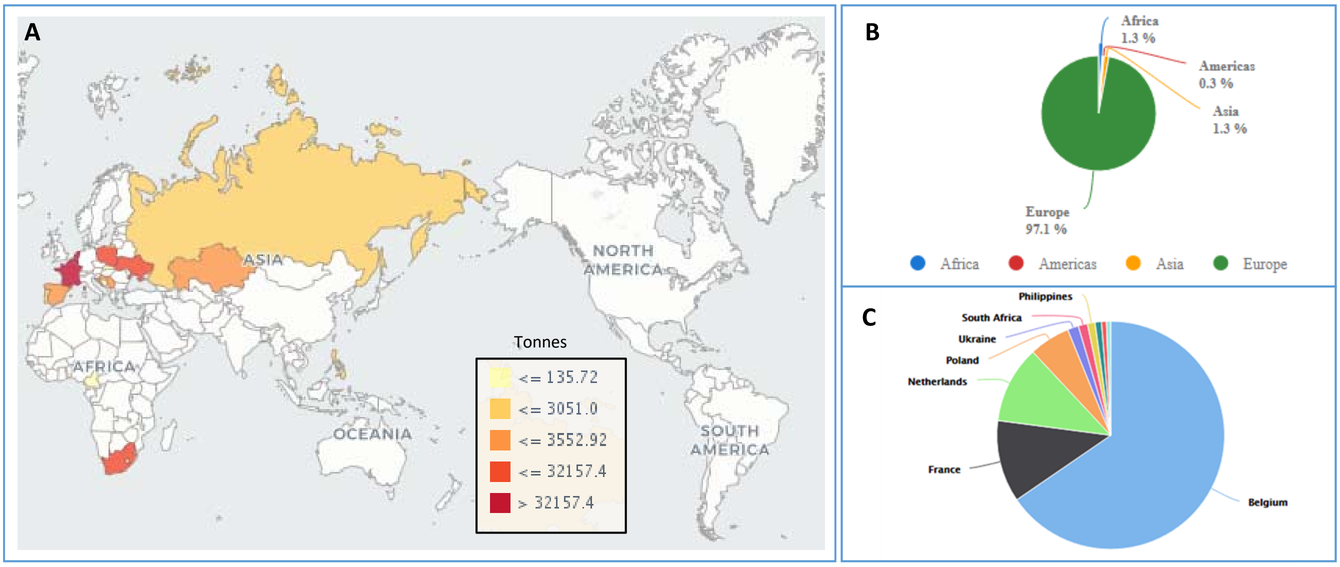


**Figure S1. Overview of worldwide chicory production.** Shown is the average over the years 1994-2018 from http://www.fao.org/faostat. A. Production quantities of Chicory roots by country, B. Production share of Chicory roots by region. C. Production share of Chicory roots by countries (top 10 contries).

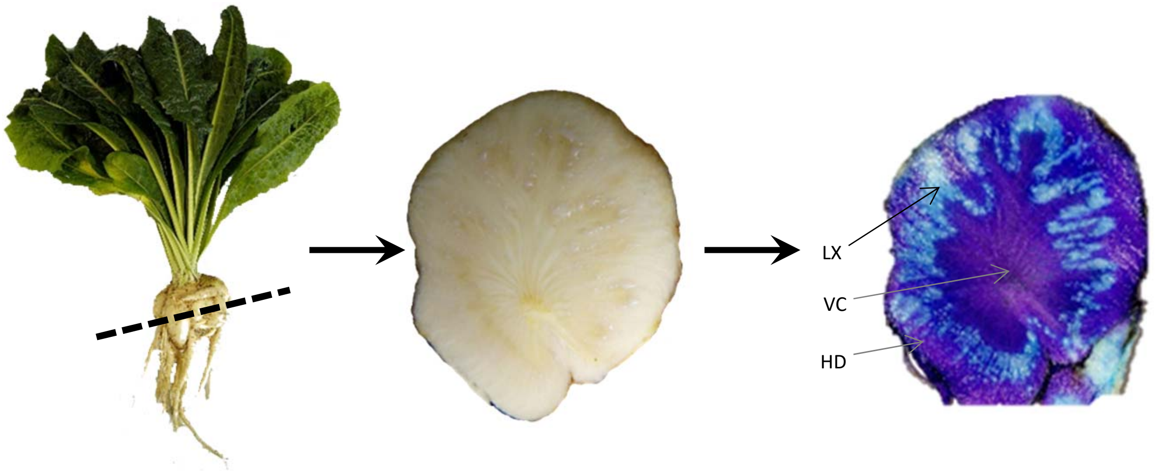


**Figure S2. Chicory root tissue visualization.** TBO staining (image on the right side) of the horizontal section of the chicory root and demonstration of the populated area with different tissues of the taproot including latex (LX), hypodermis (HD) and vascular cylinder (VC).

**A**

| | Annotated metabolite in cell types | | | annotation | | |
|---|---|---|---|---|---|---|
| | HD | LX | VC | total detected | annotated | unknown |
| GC-MS | 39 | 289 | 83 | 1657 | 330 | 1327 |
| LC-MS-Neg | 53 | 78 | 55 | 3195 | 99 | 3096 |
| LC-MS-Pos | 85 | 92 | 84 | 3631 | 112 | 3519 |
| LC-MS-hpm | 200 | 298 | 246 | 12954 | 337 | 12617 |
| total | 377 | 757 | 468 | 21437 | 878 | 20559 |

**B**

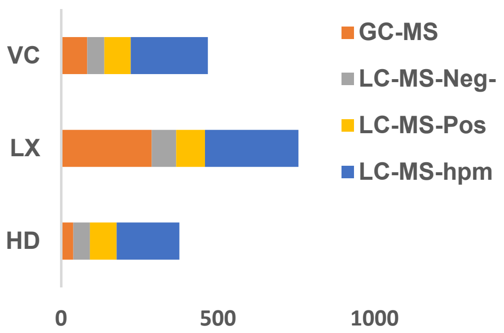


**C**

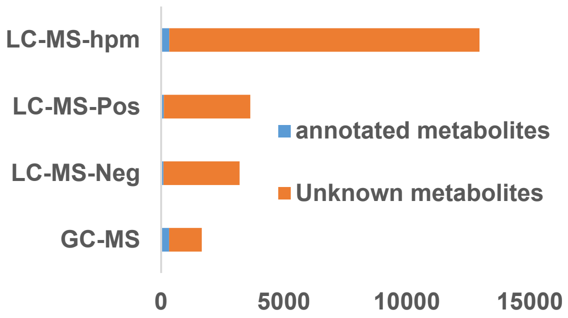


**Figure S3. Overview of the metabolomics data from chicory roots.** A. Table summarizing the number of features detected in different chicory root tissues by various MS methods and the number of annotated and unknown compounds. HD: hypodermis; LX: latex; VC: vascular cylinder. B. Graphical representation of the metabolites measured in different cell types using various MS methods. C, Graphical representation of the annotated and unknown metabolites across various MS methods. hydrophilic metabolites (hpm)

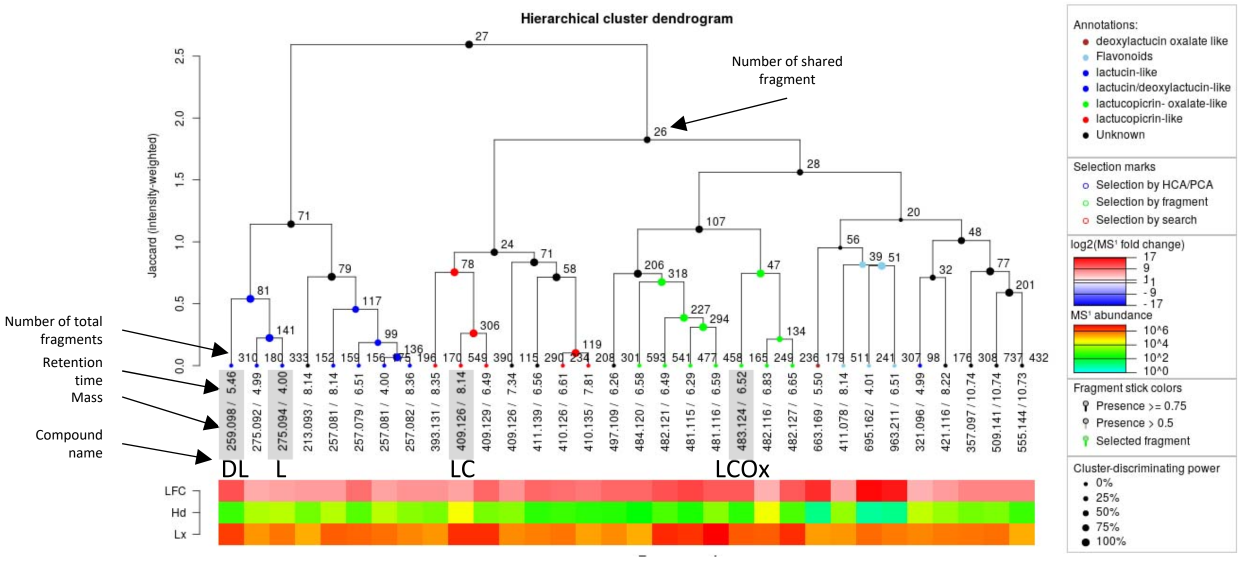


**Figure S4. Excerpt of a MetFamily output focusing on sesquiterpen lactones.** Hierarchical clustering of peak area from highly abundant fragment in LX and HD tissues measured by LC-MS-Pos. DL: deoxylactucin; L: lactucin; LC: lactucopicrin; LCOx: Lactucopicrin-oxalate.

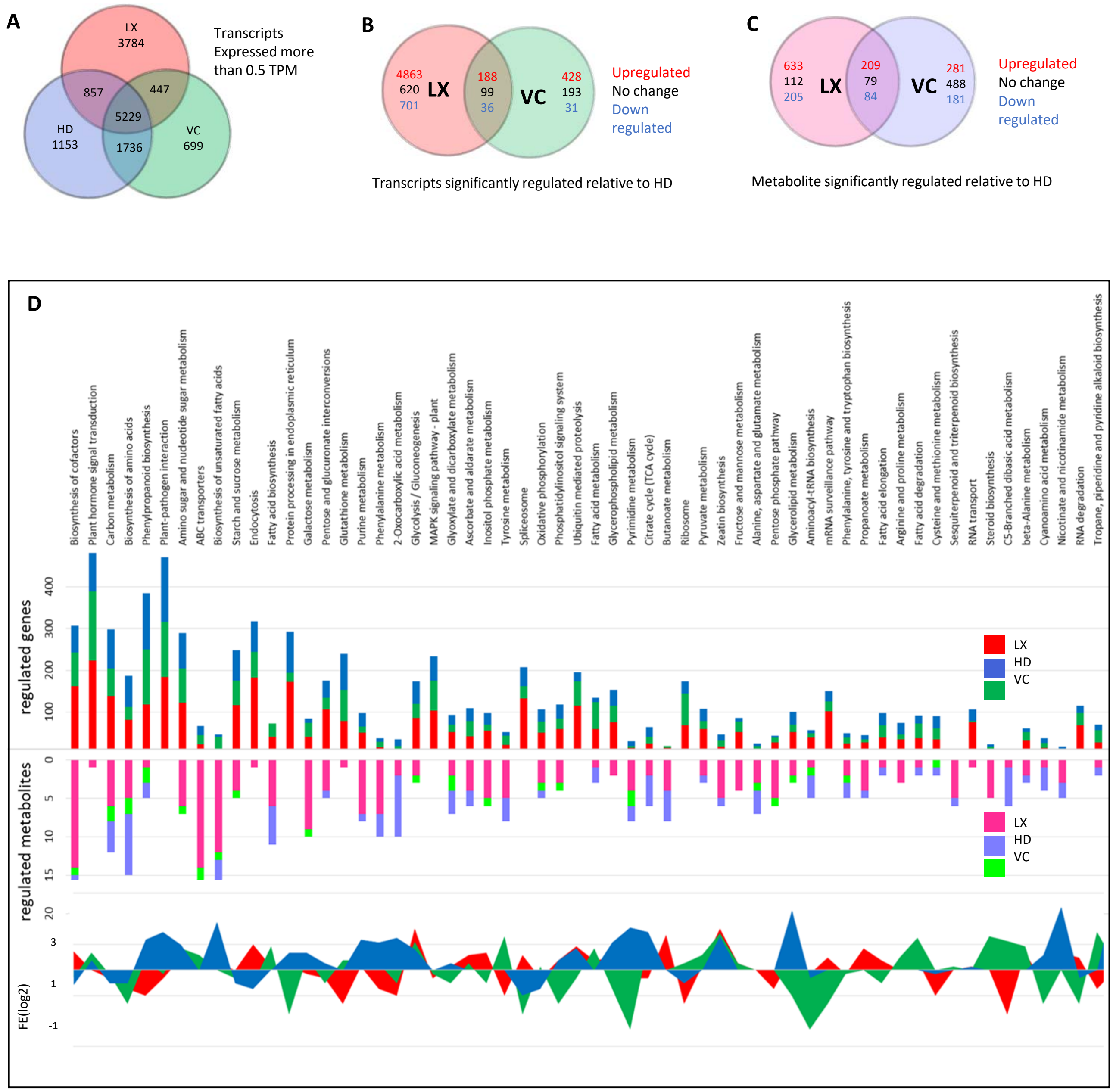


**Figure S5. Overview of transcriptomics data. A**. Frequency pattern of regulated transcripts across different cell types. Transcripts (**B**) and metabolites (**C**) that are significantly regulated relative to HD. **D.** Tissue specific transcriptome and metabolome co-analysis of the cell types in root chicory using KEGG pathways mapping and their FE (fold enrichment) to the respected pathway.

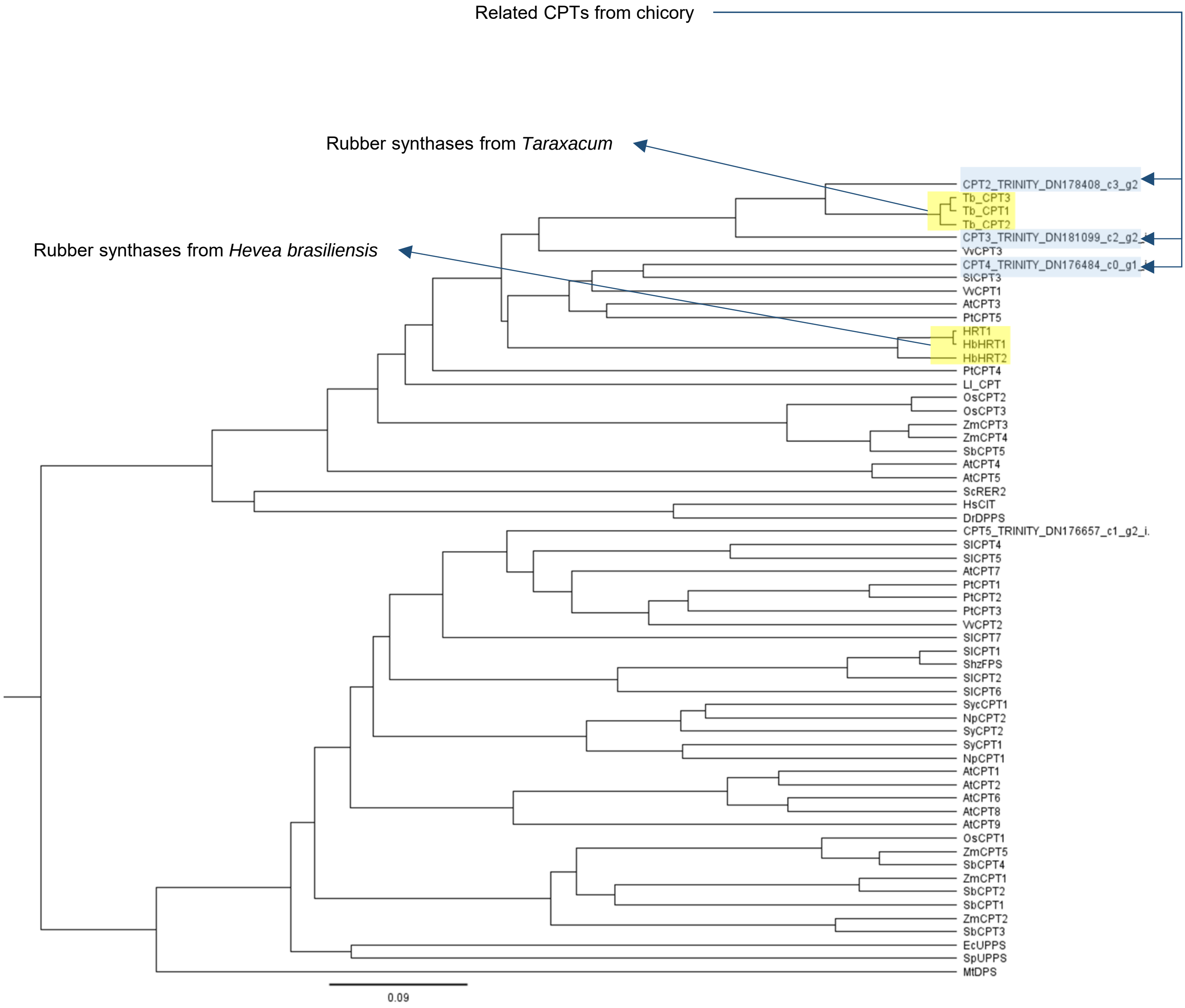


**Figure S6**. **Unrooted phylogenetic tree of cis-prenyltransferases including chicory CPT candidates and CPT genes that were described to be involved in rubber biosynthesis.** cis-isoprenyltransferase (CIT), cis-prenyltransferase (CPT), dehydrodolichyl diphosphate synthase complex (DDPS), decaprenyl diphosphate synthase (DPS), undecaprenyl pyrophosphate synthase (UPPS), z,z-farnesyl diphosphate synthase(zFPS), *Hevea brasiliensis* rubber cis-polyprenyltransferase (HRT2), *Arabidopsis thaliana* (At), *Cichorium intybus* (Ci), *Escherichia coli* (Ec), *Hevea brasiliensis* (Hb), *Homo sapiens* (Hs), *Lilium longiflorum* (Ll), *Mycobacterium tuberculosis* (Mt), *Nostoc punctiforme* (Np), *Oryza sativa* (Os), *Populus trichocarpa*(Pt), *Sorghum bicolor* (Sb), *Saccharomyces cerevisiae* (Sc), *Solanum habrochaites* (Sh), *Solanum lycopersicum* (Sl), *Streptococcus pneumoniae* (Sp), *Synechocystis* sp. (Syc), *Taraxacum brevicorniculatum* (Tb), *Vitis vinifera* (Vv) and *Zea mays* (Zm).
The tree was generated from protein sequences using Geneious v7.0 using a neighbror-joining algorithm.

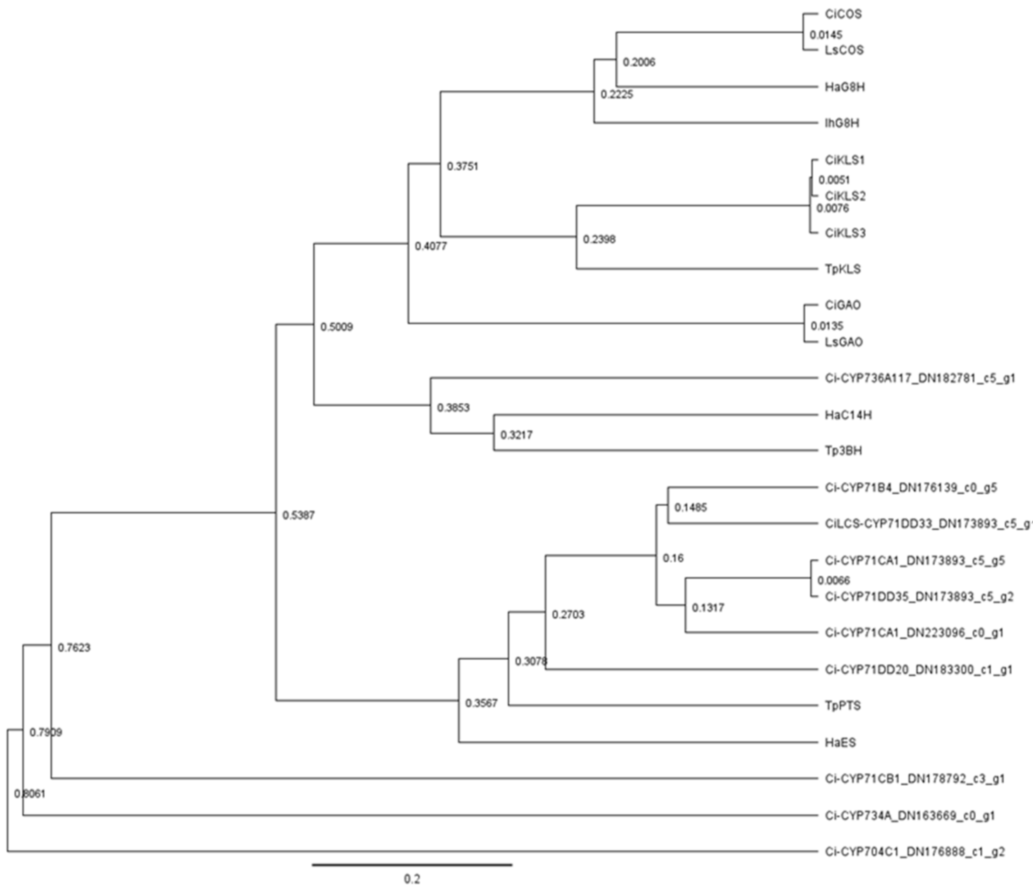


**Figure S7. Unrooted phylogenetic tree of chicory cytochrome P450 oxygenases (Ci-CYP) candidates that are overexpressed in the latex.** This includes other cytochrome P450 genes that were described to be involved in STL biosynthesis in the Asteraceae family including *Cichorium intybus* kauniolide synthase (CiKLS, ON456175), *Cichorium intybus* lactucin synthase (CiLCS, OP973199), *Lactuca sativa* germacrene A oxidase (LsGAO, GU198171), *Cichorium intybus* germacrene A oxidase (CiGAO, GU256644), *Lactuca sativa* costunolide synthase (LsCOS, HQ439599), *Cichorium intybus* costunolide synthase (CiCOS, JF816041), *Helianthus annuus* germacrene A acid 8-beta-hydroxylase (HaG8H, AEI59773), *Inula hupehensis* germacrene A acid 8-beta-hydroxylase (IhG8H, KR029572*), Tanacetum parthenium* kauniolide synthase (TPKLS, MF197558), *Tanacetum parthenium* parthenolide synthase (TpPTS, KC954155), *Tanacetum parthenium* 3-beta hydroxylase (Tp3BH, KC954153), *Helianthus annuus* eupatolide synthase (HaES, AEI59778) and *Helianthus annuus* costunolide 14-hydroxylase (HaC14H, MG765530). The tree was generated from protein sequences using Geneious v7.0 with the neighbour-joining algorithm.
